# Limited role of differential fractionation in genome content variation and function in maize (*Zea mays L.*) inbred lines

**DOI:** 10.1101/179119

**Authors:** Alex B. Brohammer, Thomas JY. Kono, Nathan M. Springer, Suzanne E. McGaugh, Candice N. Hirsch

## Abstract

Maize is a diverse paleotetraploid species with widespread presence/absence variation and copy number variation. One mechanism through which presence/absence variation can arise is differential fractionation. Fractionation refers to the loss of duplicate gene pairs from one of the maize subgenomes during diploidization and differential fractionation refers to non-shared gene loss events between individuals. We investigated the prevalence of presence/absence variation resulting from differential fractionation in the syntenic portion of the genome using two whole genome *de novo* assemblies of the inbred lines B73 and PH207. Between these two genomes, syntenic genes were highly conserved with less than 1% of syntenic genes being subject to differential fractionation. The few variable syntenic genes that were identified are unlikely to contribute to functional phenotypic variation, as there is a significant depletion of these genes in annotated gene sets. In further comparisons of 60 diverse inbred lines, non-syntenic genes were six times more likely to be variable compared to syntenic genes, suggesting that comparisons among additional genome assemblies are not likely to result in the discovery of large-scale presence/absence variation among syntenic genes.

**SIGNIFICANCE STATEMENT:** There is a large amount of presence/absence variation for gene content in maize. One mechanism that has been hypothesized to contribute to this variation is differential fractionation between individuals following the maize whole genome duplication event. Using comparative genomics, with sorghum and rice representing the ancestral state, we observed little evidence of differential fractionation among elite inbred lines and the few differentially fractionated genes identified did not appear to confer functional significance.

## INTRODUCTION

Whole-genome duplication events are prevalent throughout the lineage of many plant species. The diversification of seed plants and angiosperms occurred shortly after two whole-genome duplication events (Jiao *et al*., 2011). Whole-genome duplication events are a major driving force for angiosperm diversification and may provide a conduit for domestication to occur (Tank *et al*., 2015; Salman-Minkov *et al*., 2016). These events have profound impacts on genome structure, transcriptional regulation, biochemical functions, and ultimately phenotypes (Reviewed in PS., Soltis and DE., Soltis, 2012).

Maize (*Zea mays*) has a long history of genetic analyses that have led to the current understanding of its polyploid history, including the most recent allopolyploid event. The ancient progenitors of maize split from one another around 12 million years ago, closely following the divergence of maize and sorghum (Swigonová *et al*., 2004). The subsequent hybridization of the two maize progenitors created duplicate copies of genes genome-wide (homeologs). Synteny analysis revealed that ∼60% of maize genes are co-orthologous to a location in the ancestral state of brachypodium, rice, or sorghum (Schnable *et al*., 2012). Each homoeologous gene is derived from one of the maize progenitors and as such there exists two maize subgenomes, maize1 and maize2, that together make up the modern paleotetetraploid maize genome.

Following this most recent whole-genome duplication event, maize underwent chromosomal breakage and fusion events that recessed the 2n = 40 allotetraploid state to the 2n = 20 diploid state. This process of diploidization lead to extensive fractionation, the loss of a gene from a homoeologous gene pair, to occur in the maize genome (Langham *et al*., 2004; Woodhouse *et al*., 2010). This phenomenon of genome fractionation is distinct from other forms of DNA removal in that it is associated with the loss of genic sequence, rather than repetitive sequence, despite occurrence via the same intrachromosomal deletion mechanism (Woodhouse *et al*., 2010). Homeolog loss has been shown to be biased such that homeologs from the maize2 subgenome are 2.3 times more likely to be lost than the homeologs from the maize1 subgenome (Woodhouse *et al*., 2010). Maize also exhibits unbalanced homeolog expression bias, with the maize1 gene copy often being more highly expressed when both homeologs are retained (Schnable *et al*., 2011). Furthermore, syntenic genes and, in particular, genes from the maize1 subgenome are more likely to contribute to maize phenotypic variation (Renny-Byfield *et al*., 2017; Schnable and Freeling, 2011), and are subject to higher levels of purifying selection (Pophaly and Tellier, 2015).

Analyses within maize have shown that copy number variation (CNV) and presence/absence variation (PAV) are common throughout the genome (Springer *et al*., 2009; Swanson-Wagner *et al*., 2010; Lai *et al*., 2010; Hansey *et al*., 2012; Hirsch *et al*., 2014). There is an emerging body of evidence that indicates this genome content variation has important consequences for the extensive phenotypic variation present in maize (Chia *et al*., 2012; Maron *et al*., 2013; Hirsch *et al*., 2014; Lu *et al*., 2015). Copy number variation and PAV has been shown to underlie variation for important agronomic traits such as aluminum tolerance (Maron *et al*., 2013), starch metabolism (Haro von Mogel *et al*., 2013), flowering time (Nitcher *et al*., 2013), biochemical networks (Winzer *et al*., 2012), and disease resistance (Cook *et al*., 2012). Further, this form of variation is dispersed throughout the genome and is enriched among loci identified in genome wide association (GWAS) studies (Chia *et al*., 2012; Wallace *et al*., 2014; Lu *et al*., 2015).

Fractionation is thought to be an ongoing process (Woodhouse *et al*., 2010, Schnable *et al*., 2011), and as such can lead to continual differential fractionation between individuals within the species. Differential fractionation is one mechanism that can create PAV within individuals of a species and is defined by multiple cycles of post-tetraploidy gene loss that results in differences in syntenic gene content among individuals within a species. Maize is uniquely positioned to assess differential fraction due to the recent whole genome duplication event and the availability of two whole genome *de novo* assemblies (B73 and PH207; Jiao *et al*., 2017; Hirsch *et al*., 2016). In this study, we sought to evaluate the prevalence of differential fractionation among maize inbred lines and to assess the functional significance of this variation with regards to expression variation and phenotypic variation.

## RESULTS

### Macro-level synteny between B73 and PH207 is nearly identical

SynMap (Lyons *et al*., 2008) was used to identify blocks of syntenic genes between the maize genome (both B73 and PH207) and the sorghum and rice genome (Figure S1). On a macro scale, there were no substantial differences in syntenic block composition or subgenome assignment between the two maize genomes (Figure 1 and Figure S1). The maize1 subgenome collectively encompassed 55% (1.16 Gb) of the B73 genome and 55% (1.13 Gb) of the PH207 genome, while the maize2 subgenome encompassed 32% (0.66 Gb) of the B73 genome and 31% (0.63 Gb) of the PH207 genome.

**Figure 1.**
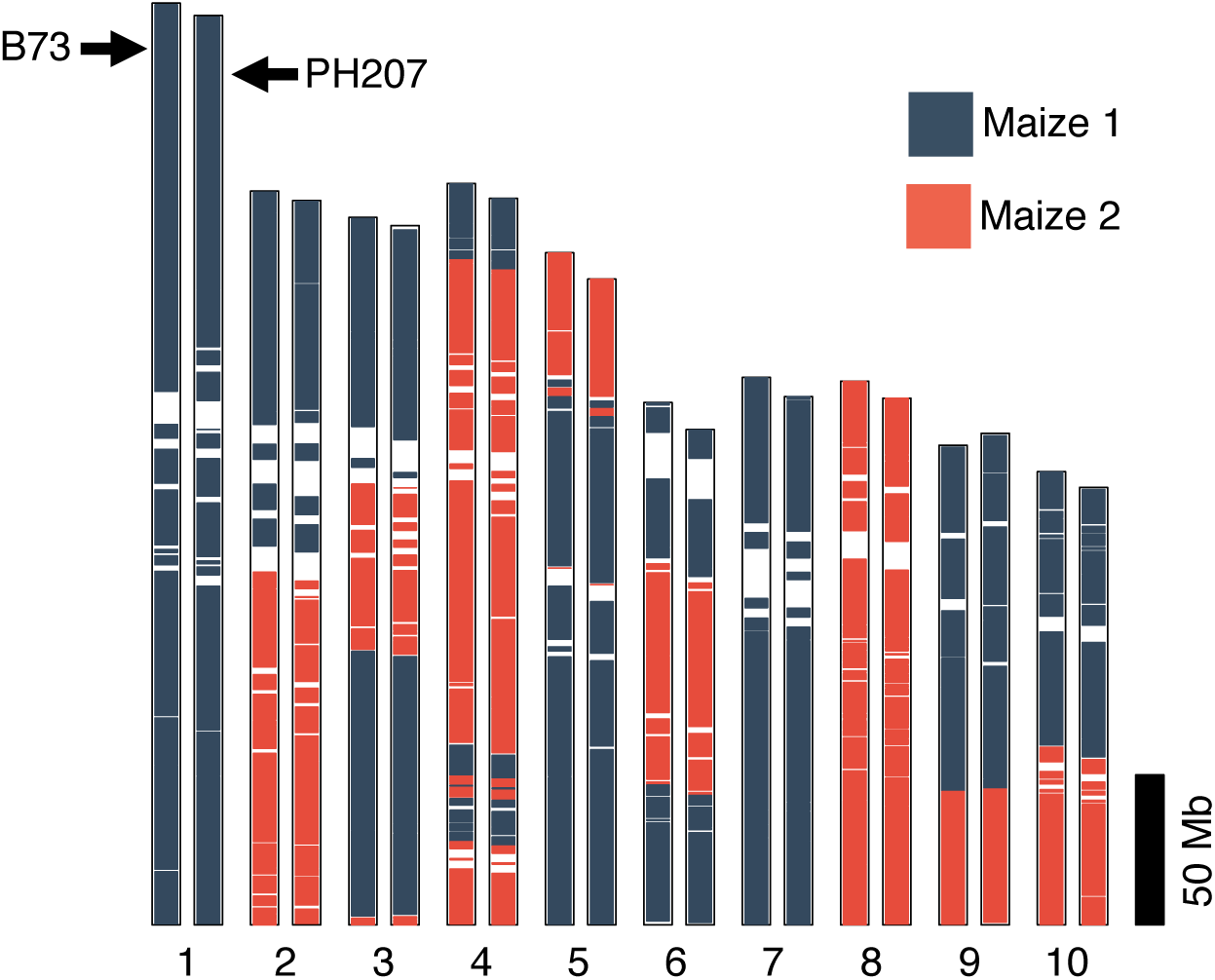
Maize subgenome syntenic blocks in the B73 and PH207 genomes. The 10 maize chromosomes are represented with B73 on the left and PH207 on the right. Syntenic blocks for the maize1 and maize2 subgenomes were determined based on comparison with the ancestral state from sorghum and rice.

Our syntenic annotation for the B73 v4 reference genome was largely consistent with reports based on current and past versions of the B73 genome assembly (Schnable *et al*., 2009; Jiao *et al*., 2017). In total, the raw SynMap analysis identified 21,568 genes in B73 and 20,446 genes in PH207 with syntenic orthologs in sorghum plus an additional 408 genes in B73 and 421 genes in PH207 that could only be identified through comparisons of rice orthologs.

### Curation of syntenic assignments

The number of syntenic genes identified through SynMap could be underestimated due to limitations of the syntenic identification software, assembly errors, or incomplete gene annotation in one or both assemblies. Significant effort was undertaken to validate and identify missed syntenic assignments including a series of BLAST alignments, alignment of resequencing data, and curating assembly gaps (see Experimental Procedures).

There were many regions in the B73 and/or PH207 genome with homology to syntenic loci that were not annotated as gene models that may reflect putative, unannotated genes. Inconsistent annotation of these genes in B73 or PH207, could lead to false positive identification of differentially fractionated genes. All putative differentially fractionated genes were aligned to the expected syntenic position in the opposite genotype (e.g. the B73 gene putatively lost in PH207 was aligned to the expected syntenic location in PH207) and mapping coordinates of each gene that could be aligned in place of an annotated gene model were included in downstream analysis. The final list of syntenic assignments included 354 loci in B73 and 1,148 loci in PH207 that likely represent bona fide gene models that were not previously annotated. In B73, 49.9% of the loci, and in PH207 55.6% of the loci had RNA-seq read coverage from at least one of five sampled tissues, providing evidence that these loci largely represent missed gene annotations.

Another major source of false negative assignments resulted from fused gene models (i.e. two separate gene models in one genotype correspond to a single “fused” gene model in another), which were identified based on significant mapping of a single gene 7 model in one genotype to two or more adjacent genes in the opposite genotype. In total, 442 instances of fused gene models in B73 and 314 instances in PH207 were identified and removed from down stream analysis.

To correct for false positive differentially fractionated genes in downstream analyses due to gaps in either of the maize genome assemblies, the putative differentially fractionated genes were aligned to annotated genes present on scaffolds or contigs and significant hits were incorporated into the working list of syntenic assignments. Putative differentially fractionated genes within gaps in the assembly were also identified and this information was included in the list of syntenic gene assignments (Table S1). This list contains 11 B73 and 219 PH207 putative gene models that could not be identified due to assembly gaps.

After the extensive curation of incorrect assignments and recovery of missing assignments the final set of maize syntenic orthologs identified in B73 included 24,514 genes and 24,454 PH207 genes (Table 1). These assignments confirm previous observations of biased fractionation between the maize1 and mazie2 subgenomes in B73 (Woodhouse *et al*., 2010), and extend this observation to a second maize genome. We identified 9,255 and 9,239 maize1 singletons in B73 and PH207 respectively, compared to 3,777 (B73) and 3,789 (PH207) maize2 singletons which supports the finding that genes from the maize2 subgenome are approximately 2.5 times more likely to fractionate than genes from the opposite subgenome (Table 1).

**Table 1.** Summary of retained duplicate, singleton, and total maize1 and maize2 genes in the B73 and PH207 genomes based on comparison to the ancestral state from sorghum and rice.

| **Gene Classification** | **B73** | **PH207** |
| --- | --- | --- |
| Retained Duplicates | 5,741 | 5,713 |
| Maize1 Singletons | 9,255 | 9,239 |
| Maize2 Singletons | 3,777 | 3,789 |
| Total maize1 genes | 14,996 | 14,952 |
| Total maize2 genes | 9,518 | 9,502 |

### Differential fractionation is not a primary driver of gene content variation

Fractionation is an ongoing process within genomes (Woodhouse *et al*., 2010), and as such can lead to differential fractionation between individuals within a species. After validating the working list of syntenic assignments and recovering missed assignments, we sought to characterize the prevalence of differential fractionation between B73 and PH207. In total, we identified 112 genes that were putatively fractionated only in B73 and 172 that were putatively fractionated only in PH207 (Figure 2A; Class II-IV). Figure 2A shows an example of a differentially fractionated gene. Of the differentially fractionated genes, 49 (B73) and 93 (PH207) were lost from the maize1 subgenome, while 63 (B73) and 79 (PH207) were lost from the maize2 subgenome.

**Figure 2.**
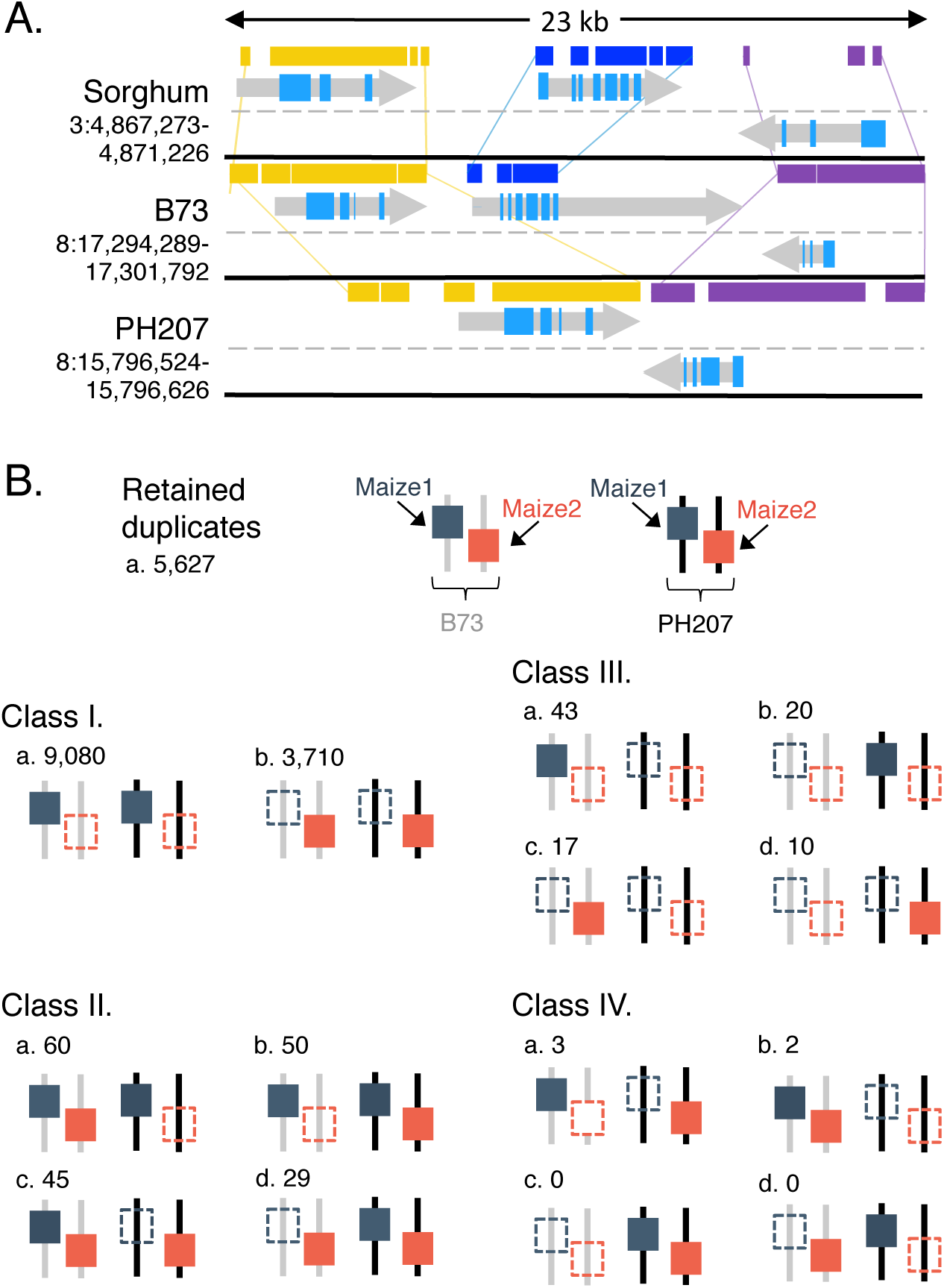
Differential gene fractionation scenarios. A) Example differentially fractionated gene. Shown are three pairs of orthologous genes in B73 and sorghum. PH207 contains only the flanking orthologous genes but is missing the homologous B73 gene, Zm00001d008692, and the sorghum gene, Sobic.003G053700. Colored blocks represent high-scoring pairs (HSPs) from alignment between each orthologous genes. B) Differentially fractionation events grouped according to the fractionation outcome observed between B73 (grey, left) and PH207 (black, right).

While there are 12 possible differential fractionation scenarios, only a subset of all possible fractionation scenarios was observed. We characterized the types and relative frequency of fractionation scenarios observed for all putative differentially fractionated genes (Figure 2B). Differential fractionation scenarios have different expectations for frequency based on the level of functional redundancy and the number of unique events that are required to derive the observed state. We hypothesized that segregation for a gene loss would occur most often if another copy was present in the other subgenome, as this redundancy would be less likely to lead to a negative fitness impact. The observed frequencies were consistent with this hypothesis. The most common differential fractionation scenario observed was the presence of a singleton in one genotype and retention of both subgenome copies in the other genotype (Figure 2B; Class II). The next most frequent differential fractionation scenario was when a singleton was retained only in one genotype and the copy from the other subgenome was fractionated in both genotypes (Figure 2B; Class III). The final scenario of differential fractionation required multiple independent loss events between the two genomes, and as expected was the least frequently observed scenario (Figure 2B; Class IV).

### Differentially fractionated genes between B73 and PH207 are more likely to exhibit PAV among diverse inbred lines than other syntenic genes

The direct comparison of syntenic gene content between B73 and PH207 revealed little PAV among syntenic genes. However, high levels of PAV may still be observed in non-syntenic genes. To further determine the PAV frequency of the differentially fractioned genes identified above within the species and to extend our analysis to the non-syntenic gene set, we resequenced 60 diverse inbred lines. All genes, including non-differentially fractionated genes, with coverage over less than 20% of the length of the gene model from 12x-65x depth resequencing data were considered significantly deleted or lost. Using this criterion for PAV, 10.1% of B73 syntenic genes and 11.3% of PH207 syntenic genes were classified as PAV among the 60 lines, while 62.5% and 58.2% of B73 and PH207 non-syntenic genes were classified as PAV (Figure 3A). On average, syntenic genes with PAV were absent across 9.1% of the inbred lines, while the subset of non-syntenic genes with PAV were absent across nearly twice as many of the inbred lines (17.2%).

**Figure 3.**
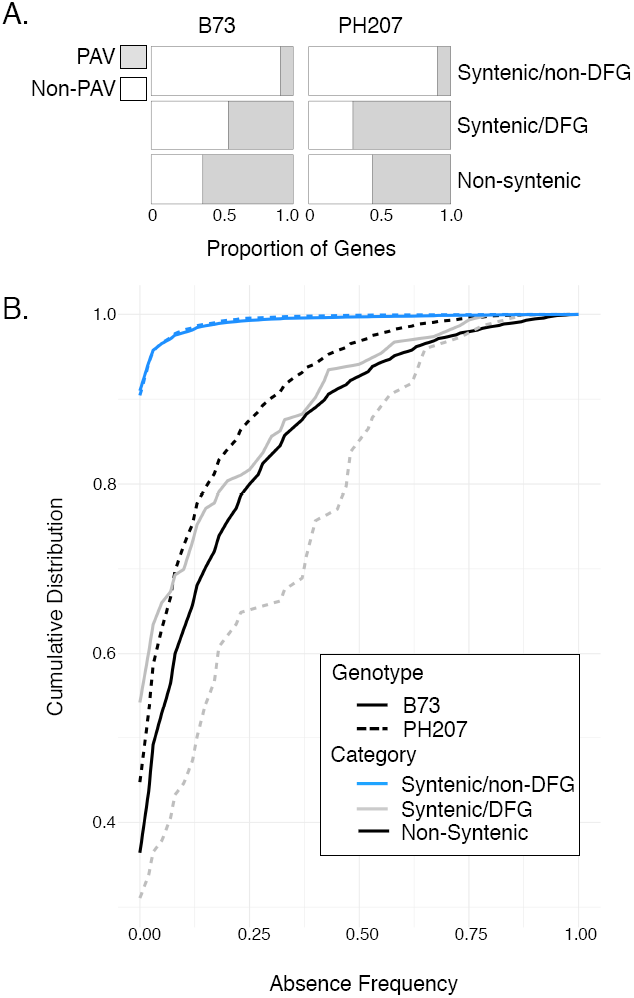
Presences/absence variation (PAV) frequency distribution of differentially fractionated genes (DFG). A) Proportion of genes that show PAV among 60 diverse lines. B) Cumulative distribution of PAV frequencies across a panel of 60 diverse inbred lines. A PAV frequency of zero indicates that a gene is not variable across any of the resequenced inbreds, while a PAV frequency of one indicates that a gene is private to B73 and/or PH207, and not contained in any of the other maize lines. Criteria for absence or substantial deletion was resequencing coverage across less than 20% of representative CDS sequence. Differentially fractionated genes are defined based on comparison of the genomes of B73 and PH207 to the ancestral state based on sorghum and rice. The distinction of B73 and PH207 in both plots is based on which genome the resequencing reads were mapped to and the subset of differentially fractionated genes that are private to either genome.

There is a clear distinction between the PAV frequency distribution of syntenic and non-syntenic genes across diverse maize inbred lines (Figure 3B). The frequency distribution for differentially fractionated syntenic genes roughly follows that of non-syntenic genes, which shows substantially higher absence frequency across diverse maize lines than syntenic non-differentially fractionated genes (Figure 3B). The deviation that is seen in the plot for the distribution of syntenic differentially fractionated genes is a result of the small number of genes in this subset of genes (112 in B73 and 172 in PH207). Additionally, the PH207 genome likely has more TEs annotated as genes than the B73 genome, causing this line to have some deviation from the distribution observed for non-syntenic genes. On average, differentially fractionated syntenic genes were absent across 16.2% of the inbreds compared to less than 1% for non-differentially fractionated syntenic genes.

### Differentially fractionated genes with a non-allelic homolog likely represent misassemblies rather than biological observations

Differentially fractionated genes that are lost from a syntenic position in the genome may have their function buffered by additional copies of the gene present in non-syntenic locations in the genome. Previous literature has suggested that the maize genome contains many homologs present in non-allelic positions throughout the genome and numerous near-identical paralogs (Liu *et al*., 2012; Emrich *et al*., 2007). Potential buffering for differentially fractionated genes was analyzed by examining coverage over non-11 syntenic gene models in resequencing data. We found that several of these genes (11/111 in B73; 43/161 in PH207), could be uniquely mapped back to a single gene model elsewhere in the genome (Table S2). Nearly all the PH207 genes (37/43) that mapped back to a non-syntenic position were located on a non-syntenic chromosome, and 2 of the 11 B73 cases mapped to a non-syntenic chromosome. The remaining B73 genes mapped to the expected chromosome but outside the syntenic block.

To determine if these non-allelic homologs were shared with any other genotypes, the draft assemblies of maize inbred lines W22 (GenBank assembly accession GCA_001644905.2), CML247 (Maize Genetics and Genomics Database, 2017), F7 (Unterseer *et al*., 2017) and Ep1 (Unterseer *et al*., 2017) were analyzed. All of the cases in which PH207 contained a non-allelic homolog on a non-syntenic chromosome were private to PH207 and retained at the B73 position in the other genotypes. Similarly, the two B73 genes with non-allelic homologs on unexpected chromosomes were only observed in B73 and not in any of the other assemblies including the previous version of the B73 assembly (Figure 4). Although we did not rule out that these are biological, evidence from multiple genomes suggests that these are predominantly the product of missassembly. These cases of potential buffering were excluded from further analysis of functional properties of differentially fractionated genes.

**Figure 4.**
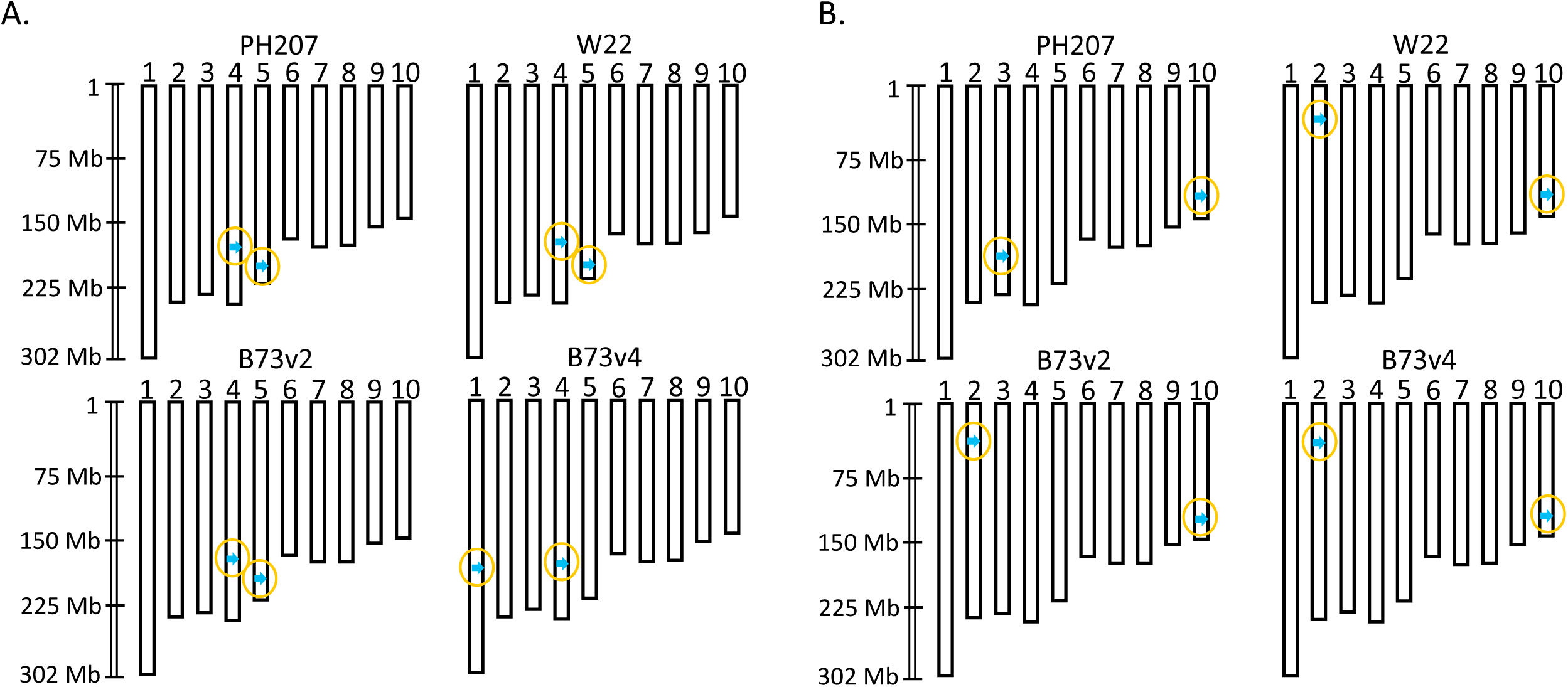
Examples of potential buffering loci from non-allelic homologs. A) The B73 gene Zm00001d031670 is present on chromosome 1 in the B73 v4 assembly (blue arrow), and present on chromosome 5 in all other assemblies (PH207, W22, CML247, F7, Ep1). The homeolog, Zm00001d052213, is present on chromosome 4 in all assemblies. B) The PH207 gene Zm00008a013424 is present on chromosome 3 in the PH207 assembly but present on chromosome 2 in all other assemblies. The homeolog of this gene, Zm00008a038016, is present on chromosome 10 in all assemblies.

### Differentially fractionated genes are underrepresented among genes of functional significance

The function of genome content variants (CNV and PAV) is generally not well understood and the specific contribution of differential fractionation to phenotypic variation has not been previously studied. To evaluate the functional consequences of differential fractionation to phenotypic variation, we tested whether differentially fractionated genes were enriched or depleted as compared to non-differentially fractionated genes in various gene sets that would indicate importance to phenotypic variation.

The first of these annotated gene sets was the maize classical gene set, which consists of 424 annotated B73 gene models that have been extensively cited in the literature and have previously been shown to be enriched for the presence of syntenic genes (Schnable and Freeling, 2011). An expanded set of 4,461 named genes manually curated by the Maize Genetics and Genomics Database (MaizeGDB.org) was combined with the classical gene list. Of the 4,649 genes in the combined list of non-redundant curated genes, 3,989 were in our list of syntenic genes. There was a significant depletion of differentially fractionated genes amongst these curated genes (Chi-square with Yates correction, one-tailed p-value 0.0274) with only 16 differentially fractionated genes overlapping with this gene set (Table 2).

**Table 2.** Overlap of differentially fractionated genes and functional maize gene lists.

| **Dataset** | **Reference** | **Overlapping**<br>**Syntenic**<br>**Genes** | **Overlapping**<br>**Differentially**<br>**Fractionated**<br>**Genes** | **Chi-Square**<br>**Test P-value** |
| --- | --- | --- | --- | --- |
| Curated Genes | (Schnable and Freeling,<br>2011; MaizeGDB.org) | 3,989 | 16 | 0.0274 |
| Hub Genes | (Walley et al., 2016) | 4,280 | 11 | 0.0005 |
| GWAS Hits | (Wallace et al., 2014) | 6,537 | 39 | 0.3735 |

The next set of genes that were tested was a set of highly interconnected ‘hub-genes’ from mRNA and protein based regulatory networks (Walley *et al*., 2016). Among the 4,280 hub-genes overlapping syntenic genes, only 11 were differentially fractionated, which was a highly significant depletion of hub-genes among the set of differentially fractionated genes (Chi-square with Yates correction, one-tailed p-value 0.0005; Table 2).

Finally, we tested a list of curated maize NAM-GWAS hits across 41 agronomically relevant traits (Wallace *et al*., 2014). As with the previous gene sets, there were few differentially fractionated genes overlapping GWAS hits, and less than 1% of GWAS hits were differentially fractionated. However, the proportion of differentially fractionated GWAS hits was not significantly different from the proportion of differentially fractionated genes among non-GWAS hits (Chi-square with Yates correction, one-tailed p-value 0.3735; Table 2).

### Differential fractionation is also limited at the transcriptome level

The term fractionation is most often invoked in terms of structural gene loss, however fractionation can also be considered at the transcriptome level. We hypothesize that transcriptional fractionation occurs at a higher rate than genome fractionation because gene inactivation can occur through mechanisms other than sequence deletion. To test the rate of transcriptional fractionation, we only considered genes for which both the maize1 and maize2 copies were retained in both B73 and PH207, and at least one homolog was expressed for a total of 4,498 maize1 homolog and 4,436 maize2 homolog sets. We detected 174 cases of “on/off” expression with the B73 gene active and 157 cases in which the PH207 gene was active across five distinct tissues. The rate of false positive detection of “off” genes based on these five tissues was evaluated in B73 using the maize gene atlas, a resource consisting of RNA-seq based expression values for 60 tissues across B73 (Stelpflug *et al*., 2016). Only 13 of the 174 B73 genes were confirmed as being not expressed across the larger set of tissues. Contrary to our hypothesis, the rate of transcriptional fractionation was nearly as rare as the rate of genome fractionation (∼0.001% transcriptional fractionation versus ∼0.006% genome fractionation). These results emphasize the high degree to which these genes are conserved at both the genome and transcriptome levels.

## DISCUSSION

Capturing genome diversity in the context of a species pan-genome has received much interest in maize and other plant research communities. These research efforts can be substantially improved by the availability of multiple reference genome assemblies within a species. One outcome of recent and ongoing efforts to assemble and annotate additional maize genomes will be to enable more accurate detection of PAV through direct genome comparisons. A deeper understanding of the role of dispensable maize genes and the mechanisms through which gene content variation is created is critical to broader pursuits such as synthetic biology. Here, we take advantage of the two annotated whole-genome maize assemblies that are currently available to study the impact of differential fractionation of genome content variation in maize and the functional consequences of this variation.

Using the B73 and PH207 assemblies, we found the vast majority of post-tetraploidy loss events were shared across inbred lines. Only 112 and 172 fractionation events were specific to B73 and PH207, respectively. This amount, which is likely an overestimate, indicates that differential fractionation has played a limited role in generating the extensive PAV that has been documented in maize (Springer *et al*., 2009; Swanson-Wagner *et al*., 2010; Lai *et al*., 2010; Hansey *et al*., 2012; Hirsch *et al*., 2014). B73 and PH207 are inbred lines that have undergone substantial selection and improvement for North American agricultural environments. Additional variation in syntenic gene content may be present in landraces and other diverse sets of germplasm that may have been subject to weaker or divergent selective pressures. Given that syntenic genes are highly enriched among functionally important genes (Schnable and Freeling, 2011), much of the variation in syntenic gene content, if it exists, is likely deleterious variation in the context of modern agricultural systems.

The few syntenic genes that are variable across maize breeding lines are an exception and are unlikely to underlie major phenotypes. Only a small proportion of the variable genes we identified overlap with hub-genes, community curated genes, or GWAS hits. Except for the latter, these overlaps indicate a significant depletion of differentially fractionated genes among functional gene sets. The failure to meet the significance threshold for GWAS hits may be a result of differentially fractionated genes being more likely to confer minor quantitative effects compared to genes with major qualitative importance.

Transcriptional loss can also result in phenotypic outcomes that are equally impactful as those generated through sequence loss. There are several examples of altered transcriptional patterns brought about through transposon insertions that are associated with discernible phenotypic effects (i.e. *Vgt1*, Salvi *et al*., 2007 and Tb1, Studer *et al*., 2011), and GWAS using transcript PAV as a marker has identified significant associations where there was not allelic variation (Hirsch *et al*., 2014). Due to the numerous mechanisms involved with terminating transcription of a gene (i.e. promoter disruption, methylation changes, etc.) versus sequence deletion, we hypothesized that this variation would be more common. However, we observed transcriptional fractionation at as low or lower of a rate as was observed for genome fractionation of syntenic genes. While there are some examples in the literature, and we identified a small number (<20 after accounting for false positives) of additional transcriptional fractionation events, these are not likely to be driving substantial phenotypic variation within the species.

Our goal was to assess the contribution of differential fractionation to maize genome content variation and to better understand how this variation relates to functional outcomes. Our results suggest that differential fractionation plays a minor role in the high levels of PAV in the maize genome and much of the existing PAV among syntenic genes is not likely to have major functional significance. Syntenic and non-syntenic gene sets have different evolutionary constraints and it is becoming increasingly clear that PAV distributions follow different frequencies among these classes of genes. While PAV among syntenic genes may be an “evolutionary dead-end,” due to purifying selection the same form of variation among non-syntenic genes may underlie variation for traits of agronomic importance such as environmental response and metabolite production. This variation may be particularly relevant to breeders because it is more likely to be segregating in breeding populations and can be important targets for future selection.

## EXPERIMENTAL PROCEDURES

### Syntenic gene identification

To identify maize genes in syntenic blocks relative to the ancestral state we ran the SynMap pipeline for both the B73 v4 (Yinping Jiao *et al*., 2017), and PH207 v1 (Hirsch *et al*., 2016), maize assemblies against the sorghum v3.1 (http://phytozome.jgi.doe.gov/) and rice v7 (Ouyang *et al*., 2007) genomes as the ancestral anchor states. All assemblies were downloaded from Phytozome 12.0.2 (https://phytozome.jgi.doe.gov), except for the B73 v4 assembly, which was downloaded from Gramene release-33 (http://www.gramene.org). SynMap was run using Quota Align to merge syntenic blocks with a coverage depth ratio of 1:2 and the tandem duplication distance was set to 15. All other SynMap parameters were set to default. Homologous genes between B73 and PH207 were identified by assignment to the same ancestral orthologs based first on assignment to the same sorghum syntenic ortholog and then the same rice syntenic ortholog for any maize genes that did not have a defined ancestral state from the comparison with sorghum.

### Curation of tandem duplicate genes

Clusters of genes identified as tandem duplicates were filtered to a single representative copy prior to assignment to syntenic regions in the maize genomes (Figure S2). For each group of putative tandem duplicate genes, the amino acid sequences from the longest transcript of each gene were aligned using clustal-omega (Sievers *et al*., 2011), back-translated to nucleotides using the annotated CDS of each gene, and pairwise similarity between the genes was estimated with the compute program from the analysis package (Thornton, 2003). Tandem duplicate genes that had less than 75% sequence identity or more than 50% gapped sites were considered misassigned tandem duplicates.

For correctly assigned tandem duplicates, the leftmost position gene of each cluster was chosen to represent the syntenic relationship. For the misassigned cases, syntenic relationships were inferred based on synonymous divergence from a putative ancestral gene. For each group of false tandem duplicate genes, clustal-omega (Sievers *et al*., 2011) was used to align the amino acid sequences of the longest transcripts and the amino acid sequence of the ancestral gene, as reported by SynMap (Lyons *et al*., 2008). Alignments were back-translated to nucleotide sequences, and synonymous divergence between each maize gene and the ancestral gene was estimated with the yn00 program in PAML (Yang, 2007). The maize gene that showed the lowest synonymous divergence from the ancestral gene was chosen to represent the syntenic relationship.

Orthofinder (Emms and Kelly, 2015) was used to identify additional ancestral orthologs within the misassigned tandem duplicates. Representative amino acid sequences from 13 grass species, excluding maize, were used as input for Orthofinder (*Aegilops tauschii*, ASM34733v1; *Brachypodium distachyon*, 3.1; *Hordeum vulgare*, ASM32608v1; *Leersia perrieri*, Lperr_V1.4; *Oropetium thomaeum* 1.0; *Oryza sativa*, IRGSP-1.0; *Panicum hallii*, 2.0; *Panicum virgatum*, 1.1; *Phyllostachys edulis*, 1.0; *Setaria italica*, 2.2; *Sorghum bicolor* 3.1; *Triticum aestivum*, TGACv1; *Triticum Urartu*, ASM34745v1). Amino acid sequences from B73 and PH207 were treated as originating from separate species, to allow for genotype-specific orthology inference. Orthologous relationships between maize and sorghum, and maize and rice were used to identify additional syntenic ancestral orthologs for non-representative false tandem duplicates.

### Validation and recovery of syntenic assignments

The subgenome identity of each chromosome was determined using a previously described method (Schnable *et al*., 2011). The percent of the genome in syntenic blocks was calculated by determining the order of each maize gene on its respective chromosome, scanning for consecutive runs of genes that were within 20 genes of one another, and appending the distance between each consecutive gene. If two genes were separated by more than 20 genes within a chromosome a new block was formed.

To remove false positive syntenic assignments, all possible pairwise BLASTN (Altschul *et al*., 1990) alignments of CDS sequences between homeologs within B73 and PH207 and between homologs across B73 and PH207 were made. An alignment threshold of 75% identity over at least 50% of the sequence length was used to remove any false or highly diverged assignments that were present in the raw SynMap output. Some genes may fail to meet the pairwise alignment thresholds due to inconsistencies in annotation. Genes that did not meet the threshold were realigned to the genome using BLASTN with the requirement that the gene map within 1-Mb of the original gene coordinates using the same coverage and identity criteria. Genes that did not meet the alignment criteria to the genome were filtered from the working file of syntenic assignments and replaced with NA (Table S1).

To remove false negative syntenic assignments that were not assigned in the original SynMap output, BLASTN was used to find significant alignments in collinear regions. A significant hit (E-value < 1e-30 and at least 75% identity over at least 50% of the sequence length), was classified as collinear by scanning for the nearest upstream and downstream syntenic genes on the expected chromosome, extracting the coordinates for these genes, and requiring that the hit be located within the window between those two genes. A buffer of 50-kb was added on both sides of the window to allow for local rearrangements that are biological or brought about by misassembly. The mapping coordinates of the genes that aligned to the expected syntenic positions were used as input to the intersect tool implemented in the BEDTools suite v2.25.0 (Quinlan and Hall, 2010) to determine if the mapping position corresponded to an annotated gene model. If no intersect with an annotated gene model was present the coordinates of the alignment were filled in to the syntenic assignments.

In some cases, an ancestral gene in either sorghum or rice or a maize gene in either B73 or PH207 was duplicated in our working list of syntenic assignments. In a subset of these cases a gene was ambiguously assigned to different gene models due to gene models that physically overlapped one another in the genome. To remove any incorrect assignments due to this case, the CDS sequence of all maize genes associated with the duplicated gene were extracted and aligned to the ancestral sequences associated with the duplicated gene using TBLASTX (Altschul *et al*., 1990). The true orthologous gene was chosen based on the highest alignment score. If a gene aligned significantly to two or more adjacent genes due to inconsistent annotations across genomes, the maize genes from both B73 and PH207 were excluded from subsequent analyses.

### Curation of differentially fractioned genes

Differentially fractionated genes were defined as those present in one maize genome (i.e. B73 or PH207) and absent in the other based on the synteny assignments described above (Table S1). The list of differentially fractionated genes was curated to remove false positives by first aligning all syntenic genes present only in one genome to the scaffold and contig sequences in the opposite genome using BLASTN. Genes that mapped significantly (E-value < 1e-30) to these locations were filtered from the working list of differentially fractionated genes.

Resequencing reads from both B73 and PH207 were then used to determine if reads from the genotype containing the fractionated gene could be aligned to the retained gene in the genome that had the retained copy. Contaminants in reads were identified with FastQC version 0.11.5 (Babraham Bioinformatics, https://www.bioinformatics.babraham.ac.uk/projects/fastqc/). Reads were cleaned of adapter contamination with Cutadapt version 1.13 (Martin, 2011). The sequences that were targeted for removal were the Illumina universal adapter, the index-specific adapter for each library, and any contaminating sequences identified by FastQC. Reads were then trimmed of low quality bases with sickle version 1.33 (Joshi and Fass, 2011), with a minimum length of 20bp, and a minimum mean quality score of 20. When one of the read pairs failed quality control, its mate was written into a single-end read file that was aligned separately. Cleaned single-end and paired-end reads were mapped to the B73 and PH207 reference genomes using bowtie2 version 2.3.0 (Langmead and Salzberg, 2012). The seed length was set to 12 bp, to adjust the sensitivity of the mapping to account for the average nucleotide diversity of maize. BAM alignments were cleaned of unmapped reads and sorted with SAMtools version 1.4 (Li *et al*., 2009). Duplicate reads were removed, and read groups were added with Picard version 2.9.2 (http://www.github.com/broadinstitute/picard). Coverage was then determined by calculating coverage of exon sequence from the longest transcript using BEDTools coverage v2.25.0 (Quinlan and Hall, 2010). Features with coverage over less than 20% of the CDS sequence were interpreted to represent high confidence gene losses.

False positive differentially fractionated genes could also arise from gaps in either of the genome assemblies. To correct these false positives, the coordinates of expected flanking syntenic genes were extracted. If 40% or more of the sequence space between the flanking syntenic genes was N’s, the fractionated gene was replaced with the coordinates for the flanking sequences and a flag indicating a gap (GAP:) was recorded in the synteny assignment file (Table S1). To identify smaller gap sequences, 5-kb of sequence on both sides of differentially fractionated genes was extracted and aligned to the homologous region from the opposite genotype using LAST (Kiełbasa *et al*., 2011). The amount of N’s between the aligned sequences was calculated as described previously and alignments with greater than 40% gaps were flagged as a gap (Table S1).

Resequencing data from 60 diverse inbred lines (Table S3 and Table S4) was used to assess the frequency of gene deletions. Reads were processed, aligned, and exon coverage was calculated as described above for the B73 and PH207 resequencing reads.

### Identification of non-allelic homologs

Fractionated genes that had read coverage over greater than 20% of the exon sequence during the Curation of Differentially Fractionated Genes (see above) can result from non-allelic homologs present in non-syntenic locations in the genome. To find the prevalence of non-allelic homologs, reads from the fractionated genome that mapped uniquely to the retained gene in the opposite genome (MAPQ score > 20) were extracted from the alignment file using Sambamba v0.6.6 (Tarasov *et al*., 2015), converted to fastq format with BEDtools bamtofastq v2.25.0 (Quinlan and Hall, 2010), and remapped back to the genotype of origin using Bowtie2 version 2.2.4 (Langmead and Salzberg, 2012) with default parameters. To determine whether the uniquely mapping reads corresponded to gene models, BEDtools intersect (v2.25.0; Quinlan and Hall, 2010) was used and coverage of each gene model was calculated using the method described above. Gene models with greater than 20% of the exon sequence covered were considered non-allelic homologs.

### Functional significance of differentially fractionated genes

Enrichment of differentially fractionated genes in lists of functional genes was used to assess the functional significance of differentially fractionated genes. The list of genes that were tested included the maize classical gene set and curated gene set (http://maizegdb.org/gene_center; accessed June 7, 2017; Schnable and Freeling, 2011), network hub-genes (Walley *et al*., 2016), and curated NAM-GWAS hits from Panzea (http://cbsusrv04.tc.cornell.edu/users/panzea/filegateway.aspx?category=GWASResults; accessed June 7, 2017; Wallace *et al*., 2014). These gene lists were generated from previous versions of the B73 genome assembly and gene annotation. Gene models were converted to the B73 v4 gene models using a conversion list obtained from Gramene (ftp://ftp.gramene.org/pub/gramene/CURRENT_RELEASE/data/gff3/zea_mays/gene_id_mapping_v3_to_v4/maize.v3TOv4.geneIDhistory.txt). Enrichment tests were conducted using Fisher’s exact test with Yates’ correction in R (R Core Team, 2014).

### Transcriptional variation analysis

RNAseq reads for B73 and PH207 from blade, cortical parenchyma, germinating kernel, root, and stele tissues were downloaded from the NCBI SRA accession number PRJNA258455. Three replicates were available for B73 and two replicates were available for PH207. Adapters were trimmed from the RNAseq reads using Cutadapt version 1.8.1 (Martin, 2011) with the quality cutoff option set to 20 and the minimum length option set to 50. Cleaned reads from B73 were aligned to the B73 genome assembly and the PH207 reads were aligned to the PH207 genome assembly using Bowtie2 version 2.2.4 (Langmead and Salzberg, 2012) and TopHat2 version 2.0.13 (Kim *et al*., 2013). Mapping parameters were set to defaults except for minimum intron size and maximum intron size, which were set to 5 bp and 60,000 bp, respectively. Transcript abundance values for longest CDS feature of each gene were calculated with HTSeq (version 0.7.2; (Anders *et al*., 2015) using the union mode and the non-strand-specific option. Gene models between the two genomes were linked using the syntenic assignments described above (Table S1). Since the abundance values were calculated using different CDS models between the two genomes that were of variable lengths, the counts were normalized by respective CDS lengths and corrected for library size differences (Table S5). For the binary expression analysis, a gene was considered transcriptionally inactive if it had three or fewer normalized counts averaged across reps in all tissues. Any gene that had a log2 fold change of 1.5 compared to the off cutoff, which was a count of eight reads, in at least one tissue was considered transcriptionally active.

### Code Availability

All code for this project is available on GitHub at https://github.umn.edu/broha006/maize-fractionation.

## ACKNOWLEDGEMENTS

This work was funded by the National Science Foundation (Grant IOS-1546727). The Minnesota Supercomputing Institute (MSI) at the University of Minnesota provided computational resources that contributed to this research. ABB was supported by the DuPont Pioneer Bill Kuhn Honorary Fellowship and the MnDRIVE Global Food Ventures Graduate Student Fellowship.

## SUPPORTING INFORMATION

**Figure S1**. Maize ancestral syntenic blocks in the B73 and PH207 genomes.

**Figure S2.** Analysis pipeline for resolving tandem duplicates.

**Table S1.** Syntenic subgenome assignments for B73 and PH207 genes using sorghum and rice as the ancestral state.

**Table S2.** Potential non-allelic homologs in the B73 and PH207 genomes that buffer differentially fractionated genes.

**Table S3.** Proximate presence/absence variation based on gene resequencing coverage in B73, PH207, and 60 diverse inbred lines mapped to the B73 reference genome assembly.

**Table S4.** Proximate presence/absence variation based on gene resequencing coverage in B73, PH207, and 60 diverse inbred lines mapped to the PH207 draft genome assembly.

**Table S5.** Normalized transcript abundance estimates for B73 and PH207 RNAseq reads mapped to their respective genome assemblies and linked by syntenic gene assignments.

